# Towards a Global Framework for Estimating Acclimation and Thermal Breadth that Predicts Risk from Climate Change

**DOI:** 10.1101/156026

**Authors:** Jason R Rohr, David J. Civitello, Jeremy M. Cohen, Elizabeth A. Roznik, Barry Sinervo, Anthony I. Dell

**Affiliations:** University of South Florida, Department of Integrative Biology, Tampa, FL 33620, USA.; Emory University, Department of Biology, Atlanta, GA 30322; University of California at Santa Cruz, Department of Ecology and Evolutionary Biology, Santa Cruz, CA 95064, USA.; National Great Rivers Research and Education Centre (NGRREC), Alton, IL, USA.; Department of Biology, Washington University in St. Louis, St. Louis, MO, USA.

**Keywords:** Acclimation, Amphibian declines, Global climate change, Phenotypic plasticity, Thermal biology

## Abstract

Thermal breadth, the range of body temperatures over which organisms perform well, and thermal acclimation, the ability to alter optimal performance temperature and critical thermal maximum or minimum with changing temperatures, reflect the capacity of organisms to respond to temperature variability and are thus crucial traits for coping with climate change. Although there are theoretical frameworks for predicting thermal breadths and acclimation, the predictions of these models have not been tested across taxa, latitudes, body sizes, traits, habitats, and methodological factors. Here, we address this knowledge gap using simulation modeling and empirical analyses of >2,000 acclimation strengths from >500 species using four datasets of ectotherms. After accounting for important statistical interactions, covariates, and experimental artifacts, we reveal that i) acclimation rate scales positively with body size contributing to a negative association between body size and thermal breadth across species and ii) acclimation capacity increases with body size, seasonality, and latitude (to mid-latitudinal regions) and is regularly underestimated for most organisms. Contrary to suggestions that plasticity theory and empirical work on thermal acclimation are incongruent, these findings are consistent with theory on phenotypic plasticity. We further validated our framework by demonstrating that it could predict global extinction risk to amphibian biodiversity from climate change.

---

Reversible thermal acclimation is an often beneficial change in a biological trait – such as metabolism, behavior, immunity, or the expression of heat shock proteins^1–4^ – in response to temperature variation^5–9^. For example, extended exposure to higher temperatures can cause a physiological change in an organism that increases its critical thermal maximum (*CT*_*max*_; mean temperature that causes disorganized locomotion, subjecting the individual to likely death)^10^ and optimal performance temperature (*T*_opt_), thus enhancing its tolerance to and reducing opportunity costs (lost foraging and mating opportunities) from higher temperatures^11,12^. Additionally, differential rates of acclimation have been proposed as a mechanism by which global climate change (GCC) indirectly causes population declines by altering species interactions^4,13,14^. Thus, acclimation ability has been proposed as a trait that allows species to cope with global warming and increased climate variability, two hallmarks of anthropogenic GCC^15–17^.

Much is known, unknown, and controversial regarding acclimation responses. For instance, theory suggests that organisms found in locations with high temperature variability might experience selection for greater acclimation abilities^6,21^ or thermal breadths – the range of body temperatures over which organisms perform well^18–20^ (Fig. 1). Both acclimation and thermal breadth are important because models of plasticity based on first principles^6,21,22^ suggest that organisms can exhibit plasticity in both their thermal breadths and their thermal modes, maxima, and minima. Nevertheless, researchers have suggested that, contrary to this theory, acclimation of thermal optima rarely occurs in laboratory experiments^6^ and the capacity for acclimation rarely correlates with the magnitude or predictability of thermal heterogeneity in the environment^6,20,23^. Hence, whether acclimation plasticity of thermal optima generally occurs and whether acclimation plasticity increases with temperature variability or latitude from tropical to mid-latitudinal regions remains controversial^18–20,23,24^. Additionally, body masses and temperature seasonality generally decrease toward the equator, especially for aquatic species^25,26,but^ ^see^ ^27^, and body mass is generally positively correlated with lifespan^28^. For these reasons, larger, longer-lived organisms are more likely to be exposed to extreme seasonal and interannual temperatures that likely select for acclimation than smaller, shorter-lived organisms. Finally, smaller-bodied species have higher mass-specific metabolic rates^28,29^, heat and cool faster due to their lower thermal inertia, and have fewer cells and physiological processes to adjust than larger organisms. For these reasons, theory based on first principles suggests that reversible acclimation capacities and rates might be positively and negatively correlated with body size across species, respectively^6,21,22,28^, but these patterns have never been demonstrated.

**Figure 1.**
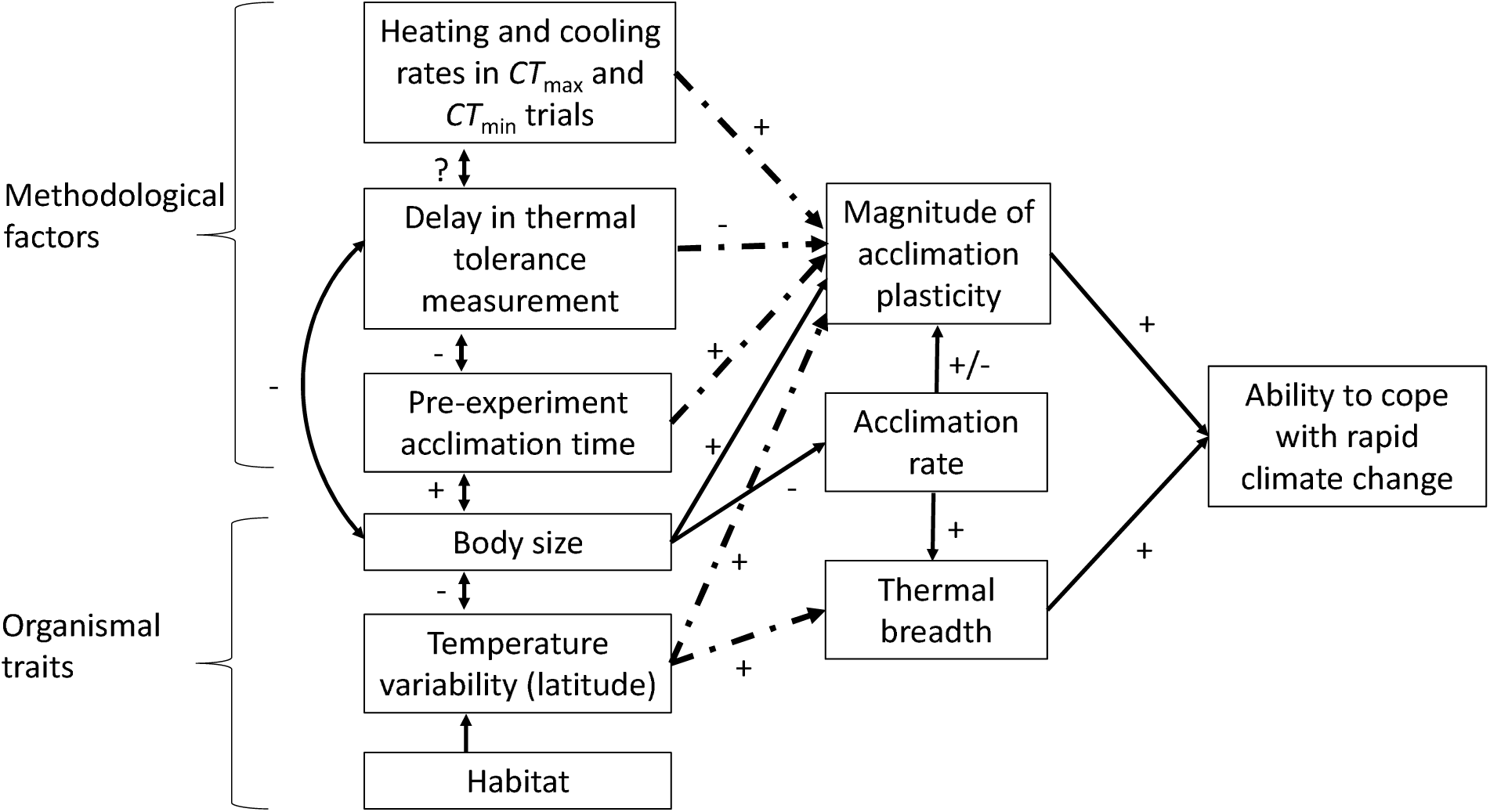
Conceptual model for how variation in methodologies and organismal traits affect the apparent ability of ectothermic organisms to cope with rapid climate change through acclimation plasticity and thermal breadth. Positive and negative signs next to arrows signify the hypothesized direction of the relationship, bidirectional arrows represent correlations because the cause-effect relationship might be unclear, and dashed arrows are effects that are hypothesized to interact statistically with organismal body size, emphasizing the considerable dependence of both acclimation and breadth on body size. This conceptual model highlights the need to more thoroughly account for methodological variation, trait variation, and their interactions to better understand acclimation and thermal breadth, as well as their relationship to coping with rapid climate change. See the primary text for a more thorough discussion of this conceptual figure.

In addition to organismal traits, acclimation responses can also be affected by experimental methodologies (Fig. 1). As an example, the strength of acclimation responses are well documented to be positively associated with acclimation duration, which is how long experimenters hold organisms at an acclimation temperature before exposing them to the test temperature^10^. This is unsurprising because acclimation takes time.

Importantly, the effects of experimental methodologies can regularly depend on organismal traits, such as body size, causing significant statistical interactions between these factors (Fig. 1); this, in turn can have several consequences for accurately measuring thermal acclimation and breadth (Fig. 1). For example, if heating rates in *CT*_*max*_ or *CT*_*min*_ trials are low, or if there is a delay between when organisms are placed at a test temperature and when trait performance is measured, then smaller organisms, because of their likely faster acclimation rates, might be more likely to acclimate to these new temperatures during trials. This will reduce the correlation between the change in acclimation temperature and the change in thermal tolerance (e.g. *CT*_*max*_, *T*_*opt*_) – a common index of acclimation abilities – resulting in a greater underestimation of the acclimation of smaller than larger organisms. Likewise, if the duration of time held at an acclimation temperature is short, there may be sufficient time for smaller but not larger species to fully acclimate, this time underestimating the acclimation abilities of larger organisms. Given the well-documented correlations among body size, latitude, temperature variability, and habitat, and experimental artifacts that can arise because of interactions between experimental methodologies and body size, biologists run the risk of drawing erroneous conclusions regarding the ability to cope with GCC unless these factors and interactions are considered simultaneously in synthetic statistical models, which is highlighted by our conceptual framework and hypotheses provided in Fig. 1. Despite the likely importance of body size to thermal acclimation, biologists understand little about how body size variation across species – or interactions among experimental methodologies, latitude, habitat, and body size –shape acclimation responses^but^ ^see19,29,30^. Once the aforementioned methodological and organismal factors are considered, we expect acclimation abilities to be greater for larger than smaller organisms, to decline from temperate to tropical regions, and for thermal breadths to be inversely correlated with body size (see SI Appendix for a discussion of how acclimation might also depend on trait identity). If these patterns emerge, they would represent the first synthesis of thermal tolerance responses to be entirely consistent with theory on thermal plasticity and metabolic rates (see^6,20,23^ for extended discussions of the inconsistency between plasticity theory and empirical results on thermal acclimation).

Here, we address these knowledge gaps and our conceptual framework and hypotheses using mathematical modeling and meta-analysis of four empirical datasets, all of which provide acclimation duration, latitude, body masses, and an index of the strength of acclimation plasticity of ectotherms (See SI Appendix, Table S1). Given that ectotherms represent ∼99.9% of all named species^31^, our analyses are relevant to most of Earth’s biodiversity (see SI Appendix for discussion on endotherms). The first dataset of Seebacher et al.^20^ provides 651 indices of acclimation strength, measured as the Q_10_ of acclimation thermal sensitivity (see Methods), for 191 species. The second dataset of Gunderson and Stillman^23^ provides 288 acclimation response ratios of *CT*_*maxs*_ for 231 species. An acclimation response ratio (ARR) is the change in a thermal tolerance measurement (e.g., *T*_*opt*_ or *CT*_*max*_) per unit change in acclimation temperature^23^ and thus, the larger the ARR the stronger the acclimation response. We added body size to both of these datasets, because neither Seebacher et al.^20^ nor Gunderson and Stillman^23^ included size in their original analyses, which we believe might explain why both studies failed to find evidence that thermal acclimation plasticity was positively associated with either temperature variability or latitude.

The third dataset we use was published by Dell et al.^32,33^ and contains 2,445 thermal response curves of ectotherms (128 of which had acclimation temperature and non-monotonic performance curves, which are necessary to compute *T*_*opt*_) measured on various traits of organisms spanning three kingdoms of life (Animalia, Fungi, and Plantae). This is also the only dataset to provide information on the thermal breadth of species (operationally defined as the width of the thermal response curve at 75% of the maximum height^6^) and the duration of time between when organisms were placed at a test temperature and when a thermal trait was first measured (See SI Appendix, Table S1). Unlike Seebacher et al.^20^ and Gunderson and Stillman^23^, the Dell et al.^32,33^ dataset runs the risk of conflating fixed and plastic responses because acclimation temperatures vary across rather than within species. However, if fixed and plastic responses were confounded, then tropical species in the Dell et al.^32,33^ dataset should have significantly warmer acclimation temperatures than temperate or polar species, but we found that acclimation temperature was uncorrelated with the absolute value of latitude (*X*^2^=0.43, *P*=0.513; controlling for habitat), suggesting that this conflation is weak at best. Nevertheless, as a precaution, we predominantly use the Dell et al.^32,33^ dataset to test hypotheses regarding thermal breadth and the duration of time between when organisms were placed at a test temperature and when performance was measured (Fig. 1). The final dataset consists of 1,040 estimates of *CT*_maxs_ for 251 species of amphibians and is used predominantly to validate our acclimation and breadth framework (Fig. 1, See SI Appendix, Table S1).

## Effect of body size on acclimation rate

We first tested the hypothesis that time to acclimate is positively related to body size (Methods). Given that acclimation takes time, the underlying assumption of these analyses is that once an organism is shifted to a new temperature, thermal tolerance will change asymptotically through time and will be faster for smaller than larger organisms. More precisely, because acclimation is a metabolic process, we predict that it should scale with body size similarly to how metabolic rate scales with mass, which scales allometrically to the one-quarter power^34,35^. Data limitations in all our datasets prohibited us from estimating a mass-scaling exponent for acclimation (see SI Appendix for details). Instead, we indirectly tested our body-size hypothesis by rationalizing that if acclimation rate is negatively correlated with size, then when acclimation duration is short, a signal of acclimation should be apparent for small but not large organisms. We found support for this hypothesis on two fronts.

First, in the Gunderson and Stillman dataset, short acclimation durations were sufficient to detect acclimation (a positive ARR) for small organisms but longer acclimation durations were necessary to detect a positive ARR for larger organisms (Three-way interaction Acc. time x mass x heat rate: *X*^2^=5.27, *P*=0.022; Fig. 2a,c). Additionally, body size and acclimation duration interacted similarly to affect acclimation signatures (i.e., a positive correlation between acclimation temperature and *T*_opt_ or *CT*_max_) in both the Dell et al. (Fig. 2b,d, See SI Appendix, Table S2) and amphibian *CT*_max_ (See SI Appendix, Table S3, Fig. S1) datasets. The minimum acclimation duration in the Seebacher et al. dataset was one week (See SI Appendix, Table S1), and thus it lacked the short acclimation periods necessary for testing effects of both short and long acclimation durations on acclimation responses.

**Figure 2.**
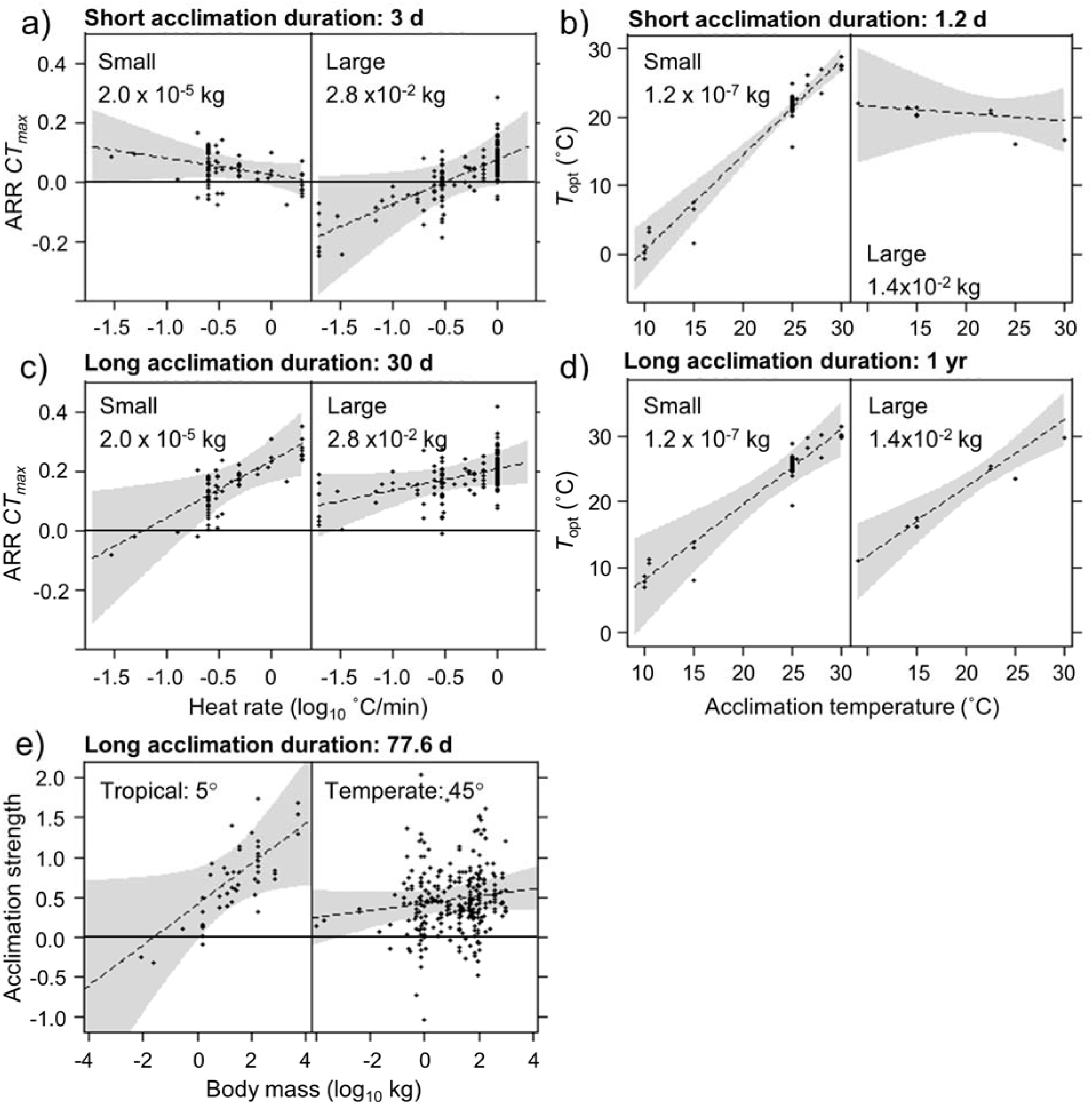
Partial residual plots showing that small organisms acclimate faster than larger organisms (a-d) and that acclimation abilities depend on an interaction between latitude and body size (e). Partial residual plots hold all factors in the statistical model that are not being displayed constant (see SI Appendix “Supplementary Discussion: Details on the visreg package”). Acclimation was measured as the acclimation response ratio (ARR) which is the correlation between acclimation temperature and critical thermal maximum (*CT*_*max*_; **a** & **c**) or optimal performance temperature (*T*_opt_; **b** & **d**). See “Methods: Statistical analyses” for the measure of acclimation strength from Seebacher et al. (2) used in **e)**. When acclimation durations are short, only smaller organisms show a positive mean ARR (**a**; i.e., they acclimate) or positive slope **b)** (see same result in See SI Appendix, Fig. S3), but when acclimation durations are long, both small and large organisms show acclimation responses (**c** & **d**; *T*_opt_ three-way interaction: *X*^2^=10.23, *P*=0.001, *n*=60; range of absolute value of latitudes 25-57°). Similarly, when acclimation durations are long, small organisms do not show positive ARRs when the heating rate in *CT*_*max*_ trials is low (presumably because they are at least partly acclimating to the new warmer temperatures during the trial), whereas large organisms show positive ARRs at most heating rates (**c**; heat rate x size x duration: *X*^2^=4,47, *P*=0.0345, *n*=262). Subpanels represent different body size categories (breaks based on 50^th^ and 90^th^ percentiles and 20^th^ and 80^th^ for *T*_opt_ and *CT*_*max*_, respectively; see SI Appendix “Supplementary Discussion: Details on the visreg package”) and short and long acclimation is represented by 20^th^ and 80^th^ percentiles. Despite smaller organisms acclimating faster than larger organisms, when acclimation durations were long (conditioned on 77.6 d, the 80^th^ percentile), large organisms showed greater acclimation capacity in general, but especially in the tropics (**e;** Latitude*mass: *z* = 2.18, *P* = 0.029). This result matches the findings from the two other datasets (see Fig. 3 and See SI Appendix, Fig. S4-5). Gray shading shows associated 95% confidence bands.

Second, the Gunderson and Stillman dataset also provided information on the heating rate of *CT*_*max*_ trials, offering another means of testing our hypothesis that time to acclimate is positively related to body size. If smaller organisms acclimate faster than larger organisms, then when the heating rate is low, smaller organisms should be more likely to partly or fully acclimate to the new warmer temperatures during the trials than larger organisms. This would reduce the ARR, thus diminishing the signal of acclimation more for smaller than larger organisms.

Consistent with this hypothesis, the Gunderson and Stillman dataset revealed that when the heating rate in *CT*_*max*_ trials was low, smaller organisms failed to show positive ARRs (confidence interval overlaps with zero on left side of Fig. 2c); in contrast, larger organisms showed positive ARRs (confidence interval almost never overlaps with zero) at most heating rates (Acc. time x mass x heat rate: *X*^2^=5.27, *P*=0.022; Fig. 2c). Hence, across a diversity of taxa, habitats, and traits, smaller organisms appear to acclimate more quickly than larger organisms.

Given that large organisms appear to take longer to fully acclimate than smaller organisms, we also tested whether the mean acclimation duration imposed by experimenters (using the Dell et al. dataset because it had the most acclimation durations) was sufficient to fully acclimate large organisms (see Methods). In these analyses, acclimation duration was independent of body size (*X*^2^=0.27, *P*=0.598), and the grand mean acclimation duration was only 85 h (or 5.49 log_10_ + 1 seconds; See SI Appendix, Fig. S2), which we show is insufficient to detect significant acclimation for most large organisms (>0.0086 kg; See SI Appendix, Table S2).

## Acclimation, body size, and latitude

Given that these initial analyses made it clear that acclimation depends on body size, acclimation rate, and acclimation duration and first principles suggest that selection for plasticity should depend on environmental varaiability^21^, we developed a mathematical model for acclimaton and thermal breadth based on the following assumptions: i) the magnitude of acclimation depends on an organism’s acclimation rate and duration up to some physiological limit, which increases with latitude^18–20^ (also see below), ii) acclimation rate scales logarithmically with body size and temperature^28^ (Fig. 2, See SI Appendix, Table S2 & 3, Fig. S3), and iii) there is a delay between when organisms are placed at a test temperature and when a trait is measured. (Methods). Our model is an extension of the seminal theoretical work of Gabriel and colleagues^21,22^ that explored how organisms shift their modes and breadths of environmental tolerance functions in response to variability in and response lags to environmental stressors. Specifically, unlike the work of Gabriel and colleagues, our model addresses the consequences of body size- and latitude-dependent rates and limits to thermal plasticity on the expression of thermal acclimation and breadth.

Our goal of the modeling exercise was to evaluate whether the mathematical model with only the assumptions above could recreate the salient relationships among acclimation duration, body size, and latitude observed in the empirical data on acclimation strength and thermal breadth. If it could, then the principle of parsimony would suggest that seminal theories on plasticity^21,22^ and the metabolic theory of ecology^28,34,35^ might indeed accurately predict true thermal tolerance responses, despite assertions to the contrary^6^. If the model could not recreate the salient relationships, then it would suggest that thermal plasticity theory was missing something, as suggested by Angilletta^6^. Although it would be ideal to develop a more sophisticated model that could generate quantitative predictions that were regressed against observed data, this was outside the scope of the current work.

Statistical analyses of our emprical data and our modeling simulations independently showed that acclimation plasticity declined from mid-latitudes to the tropics as predicted. In the Seebacher et al. dataset, significant acclimation was detectable for both small and large organisms at mid-latitudes, but only for large organisms at low latitudes (Fig. 2e, See SI Appendix, Table S5a,b). Similar patterns were apparent in the Dell et al. (Fig. 3d,e, See SI Appendix, Table S4) and amphibian *CT*_max_ (See SI Appendix, Table S6, Fig. S4) datasets. The Gunderson and Stillman dataset also showed the same pattern, although latitude was replaced by seasonality (standard deviation of annual mean weekly air temperatures; See SI Appendix, Table S7, Fig. S5), providing support for the hypothesis that the greater capacity to acclimate at midlatitudes is a function of greater variability in environmental temperature. Thus, despite smaller organisms acclimating faster than larger organisms, when acclimation durations are sufficiently long, larger organisms showed greater acclimation capacity in general, especially in the tropics where temperature variability is low relative to temperate regions. This result is consistent with theory that suggests that acclimation capacity should be correlated positively with temperature variability^21,22^ because, at a given latitude, larger organisms should experience greater temperature variation over their lifetime than smaller organisms because of their generally longer life spans. Habitat generally did not significantly interact with most predictors, and there were no consistent effects of habitat on acclimation responses across the datasets (See SI Appendix, Tables S2-S8).

**Figure 3.**
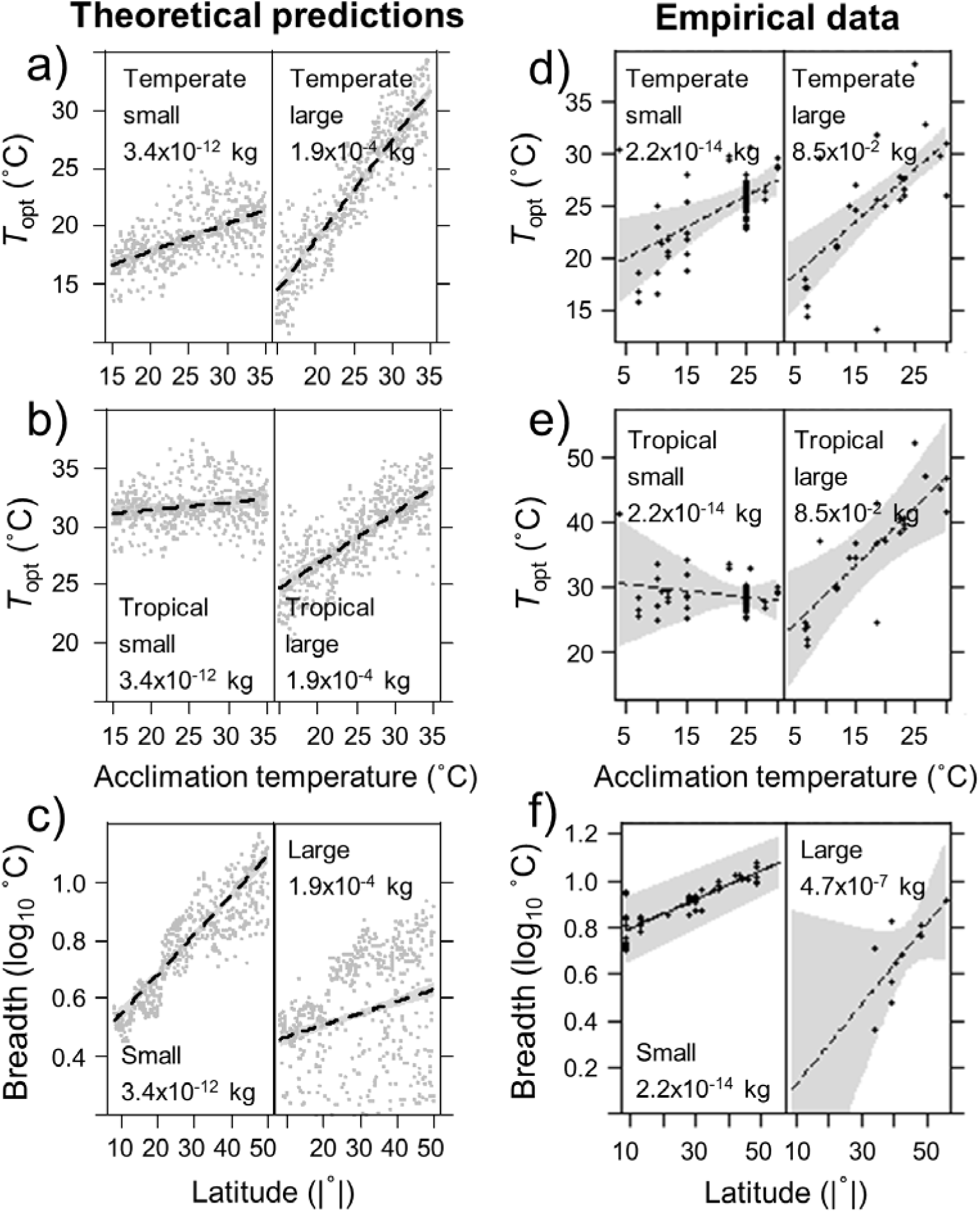
Partial residual plots showing the predicted and observed effects of acclimation temperature, body mass, and latitude on optimal performance temperature and thermal breadth for a diversity of taxa and habitats. a), b), and **c)** show results from our mathematical model for optimal performance temperature (*T*_opt_) at 45 and 5 degrees latitude and for thermal breadth, respectively (see SI Appendix for parameters). **d)**, **e)**, and **f)** show the same plots, respectively, but for Johnson-Lewin model fits (see See SI Appendix, Fig. S7 and S8 for similar results using Weibull fits) to empirical data obtained from Dell et al. dataset (three-way interaction for *T*_opt_: *X*^2^=8.08, *P=*0.0045, *n*=105; two-way interaction for breadth: *X*^2^=13.61, *P<*0.001, *n*=64; log masses <10^-5^ kg). Subpanels represent different body size categories (breaks based on 20^th^ and 80^th^ percentiles). Gray shading shows associated 95% confidence bands. See See SI Appendix, Fig. S11 for similar results from the model when no relationship between acclimation rate and temperature is assumed (assumption here is an exponential relationship).

Although we used the Seebacher et al.^20^ and Gunderson and Stillman^23^ datasets to identify latitudinal and seasonality gradients in acclimation, the original authors failed to detect such patterns. There are likely several reasons why this occured, such as not including body mass data in their analyses, which strongly interacts with both acclimation duration and latitude/seasonality, and by not weighting their analyses by sample size or variance (see SI Appendix for additional details). As we show in our conceptual framework (Fig. 1), it is important to control for methodological variation, trait variation, and their interactions, as well as correlations among organismal traits, to reliably detect the generally positive association between environmental variability and acclimation. However, a failure to control for these factors cannot explain all cases where environmental variability is unrelated to thermal acclimation^24^, and thus there are exceptions where other factors, such as phylogenetic inertia or epistasis^6^, might place limits on thermal plasticity for some species.

Our simulation model suggests that, in addition to weaker selection for acclimation in the less thermally variable tropics^18–20^, the apparent weaker acclimation of smaller organisms is a product of them acclimating so fast that much of their acclimation occurs during the delay between when they first experience the test temperature and when researchers begin measuring performance (i.e., an experimental artifact; Fig. 1). This was also supported by experimental data. Based on the entire Dell et al. dataset (1,480 curves with necessary data for analyses), body size was associated negatively with acclimation duration (*F*_1,1478_ = 41.92, *P* < 0.001, See SI Appendix, Fig. S6), a methodological pattern that can exaggerate this artifact. For example, very small organisms, such as microbes, were held at a test temperature for a mean of 8.82 h (the y-intercept) before a trait was first measured (Fig. 1), which, according to our analyses on time to acclimate (see Fig. 2), is sufficient time for substantial if not full acclimation for such small organisms.

## Thermal breadth, body size, and latitude

As with acclimation, both the simulation model and statistical analyses (See SI Appendix, Table S8) revealed consistent results for thermal breadth. Smaller organisms had greater breadths than larger organisms (Fig. 3c, f), although this difference was larger at low than mid-latitudes, at least partly because few large, tropical organisms were tested (Fig. 3f). Species exhibited an increase in thermal breadth with increasing latitude (latitude x body mass: *X*^2^=13.61, *P*<0.001; Fig. 3c, f, See SI Appendix, Table S8), confirming previous results^36^. Our model suggests that smaller organisms could appear to have greater thermal breadths than larger organisms because they acclimate more rapidly, maintaining higher observed performances over a larger range of temperatures (Fig 1, Fig. 4), although fixed responses could also explain some of this pattern. Additionally, the model highlights that the greater magnitude of acclimation that occurs at higher latitudes is a possible driver of the positive relationship between thermal breadths and latitude (Fig. 3c, f)^36^. Importantly, these acclimation and breadth results were robust to whether symmetric or asymmetric curves were used in the mathematical model (See SI Appendix) and whether Johnson-Lewin or Weibull models were fit to the thermal performance curve data (Fig. 3,4 vs SI Appendix, Fig. S7, S8).

**Figure 4.**
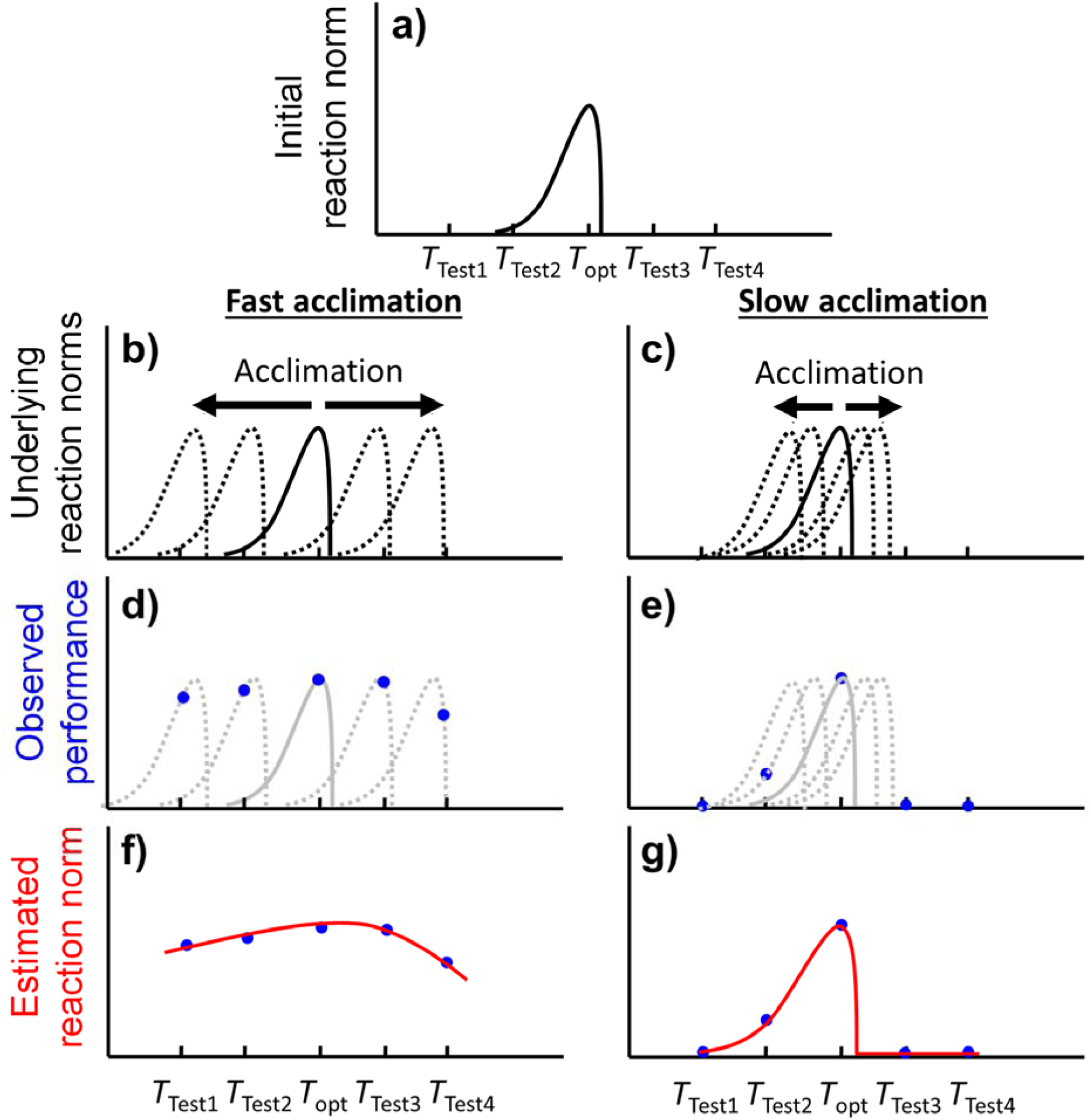
Conceptual framework connecting time to acclimate with thermal performance breadth. If organisms **a)** have thermal response curves of fixed shapes with an optimal temperature (*T*_opt_), but can acclimate either **b)** rapidly or **c)** slowly to different test temperatures (*T*_Test1_*…T*_Test4_) by sliding these reaction norms along the temperature axis during a finite acclimation time (dashed curves, one corresponding to each test temperature), then organisms that acclimate rapidly can **d)** maintain high observed performance (blue points) over a larger temperature range than **e)** those that acclimate slowly. When thermal performance curves (red lines) are fit to the resulting data, organisms that acclimate rapidly appear to have larger breadths than organisms that acclimate more slowly because they exhibit greater acclimation in the delay between when they first experience the test temperature and when researchers begin their performance measurements **f), g)**.

## Thermal plasticity, global climate change, and conservation

Our findings also help identify species that might be at risk from GCC (Fig. 1). For instance, owing to their narrower breadths and longer times to acclimate and evolve adaptations (Fig. 2,3), our model suggests that larger tropical ectotherms might experience greater lethal and sublethal effects from GCC than smaller temperate ectotherms. To test these predictions and thus further validate our model, we used the amphibian *CT*_*max*_ dataset to quantify the relationship between the thermal safety margin (*CT*_*max*_ minus maximum temperature of warmest month at location of collection)^36,37^ of 185 amphibian species – the most threatened vertebrate taxon on the planet^4^ – and their IUCN (International Union for the Conservation of Nature) threat status, controlling for various additional factors (see Methods). As our model predicted, large tropical amphibian species (with small geographic ranges; species with large ranges were rarely threatened regardless of body size or latitude, See SI Appendix, Fig. S9) had the strongest negative relationship between threat status and thermal safety margin and thus are indeed most threatened by GCC (Fig. 5a,b). Also, as predicted, this threat level decreased as latitude increased or body size decreased (interaction: *X*^2^=8.66, *P*=0.0033; Fig. 5a,b). Importantly, this relationship between threat status and thermal safety margin was detectable despite the many factors other than GCC contributing to amphibian declines^14,38,39^.

**Figure 5.**
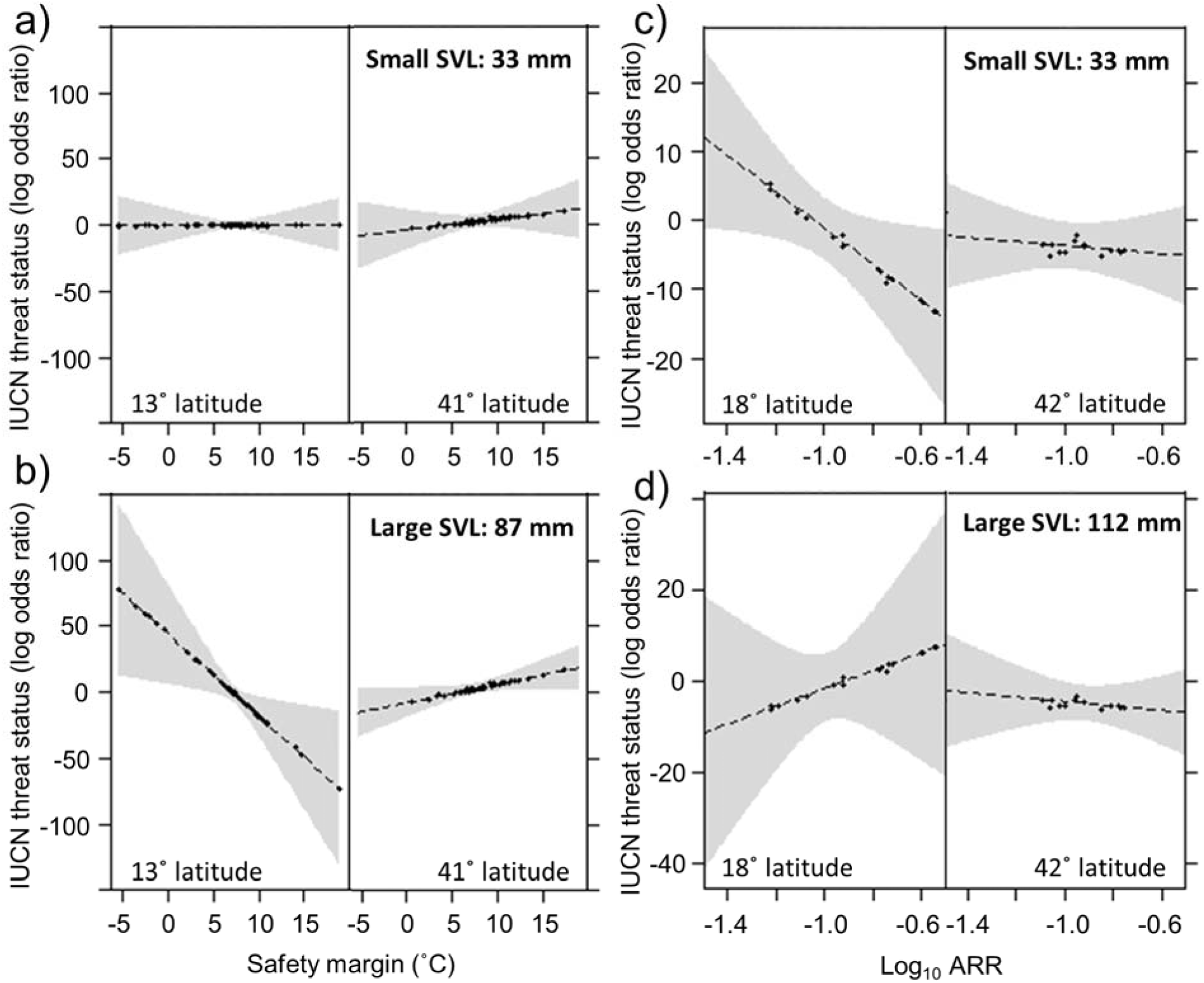
Partial residual plots showing that large tropical amphibians are most threatened by climate change at least partially because of limited acclimation abilities. **a)** and **b)** show the interaction (*X*^2^=8.66, *P=*0.0033) among thermal safety margin (critical thermal maximum – maximum temperature of warmest month), the absolute value of latitude, and log body size (**a)** small snout-vent lengths [SVL], **b)** large SVL; 20th and 80^th^ percentiles) on the odds that amphibian species with small geographic ranges (conditioned to the 20^th^ percentile; 3.8 log km^2^) are categorized as threatened or not by the IUCN (*n*=186; see See SI Appendix, Fig. S9 for species with large geographic ranges). **c)** and **d)** show the interaction (*X*^2^=8.66, *P=*0.0033) among acclimation response ratio (ARR, a measure of thermal acclimation plasticity), the absolute value of latitude, and log body size (**c)** small SVL, **d)** large SVL) on the odds that amphibian species with a small thermal safety margin (conditioned to the 20^th^ percentile; 5.8° C) are threatened (*P<*0.05; *n*=74; see See SI Appendix, Table S9 for full statistical model). Subpanels represent different latitude categories (breaks at 10^th^ and 90^th^ percentiles), revealing that large tropical amphibians are at the greatest threat from climate change (i.e., when the safety margin is small, **a**, **b**) perhaps because plasticity reduces this threat less so for large than small amphibians at low latitudes (**c**,**d**). Gray shading shows associated 95% confidence bands.

This analysis, however, does not specifically evaluate the contribution of acclimation plasticity to threat status. To do this, we re-analyzed the same dataset using the subset of amphibian species for which we also had thermal acclimation plasticity data (ARR; 74 species), testing for interactions among safety margin, ARR, body size, and latitude, and controlling for the local level of climate change (See SI Appendix, Table S9). Not surprisingly, acclimation plasticity reduced threat status the most (i.e., was the most protective) when the safety margin was small (Fig. 5c,d), highlighting the protective nature of acclimation plasticity to GCC. In fact, the slope between ARR and threat status was never significantly negative at large safety margins (See SI Appendix, Table S9). The biggest difference in threat status between large and small amphibians occurred at low latitudes (Fig. 5a,b). If differential plasticity contributed to this threat status pattern, then the greatest difference between large and small amphibians in the protectiveness of plasticity against GCC (i.e., slope between threat status and ARR) should also occur at low latitudes. The relationship between threat status and ARR became more negative as body size decreased and was similar for large and small amphibians in temperate regions.

However, the same ARR at low latitudes reduced the threat status (was less protective) of larger amphibians less than smaller ones (Fig. 5c,d); and thus, as predicted, the greatest difference between large and small amphibians in the protectiveness of plasticity against GCC occurred at low latitudes. This is likely because most *CT*_*max*_ studies ignore time to acclimate. Because smaller organisms acclimate sooner than larger organisms, even with the same ARRs, smaller organisms pay the costs of their physiology mismatching their environment for a shorter period of time. Overall, these results suggest that variation in thermal acclimation abilities might partly account for why amphibians experience a greater threat from GCC as body size increases and latitude decreases.

## Conclusions

Here we demonstrate that methodological factors, body mass, and latitude interact to shape the actual and perceived thermal acclimation responses of ectotherms. Our relatively simple mathematical model with only a few assumptions recreated the complex patterns of acclimation observed in four independent and diverse datasets consisting of experiments conducted across acclimation durations, body masses, habitats, traits, latitudes, and >500 species. Although we were unable to test the hypothesis that acclimation rate should scale with body size to the ¼ power, our model assumed this and its output was consistent with the extensive experimental data, findings congruent with the metabolic theory of ecology^28,34,35^. Additionally, our model and the experimental data suggest that the shorter times to acclimate of smaller than larger organisms drive the generally observed wider thermal breadths of smaller organisms (Fig. 4). Importantly, our findings are consistent across all four datasets, despite various strengths and limitations of each dataset. One of these datasets contained thermal optima, providing evidence that thermal optima seem to regularly acclimate despite assertions to the contrary^6^. Although other factors undoubtedly affect thermal acclimation and breadth, our results suggest we are capturing many of the principal mechanisms driving variation in acclimation and breadth across the globe and species (but see caveats, including discussion of cold acclimation, in SI Appendix). By demonstrating that acclimation abilities are greater for larger than smaller organisms and decline from temperate to tropical regions, that thermal breadths are inversely correlated with body size, and that much of the previous controversy^6,8,20,23^ regarding these relationships was a product of insufficiently accounting for methodological factors and important statistical interactions, we believe that we have offered the first synthesis of thermal tolerance responses to be entirely consistent with theory on thermal plasticity and metabolic rates. Additionally, given that body mass is strongly correlated with generation time and latitude is strongly correlated with diel variation, our findings have potential to be extended to these other common predictors of thermal acclimation^40^.

Our model identifies large tropical ectothermic species at particular risk from GCC, which was validated by evidence that large tropical amphibians already experience a greater threat from GCC than any other tested amphibian group. Our assertion that tropical ectothermic species should be more sensitive to GCC than temperate species is consistent with previous studies^36^, and our assertion that larger organisms should be more sensitive to GCC than smaller organisms is consistent with GCC reducing the body sizes of aquatic organisms^26,31^, temperature variability benefiting pathogens (small-bodied) more so than hosts (large-bodied), and recent disease emergences being linked to GCC^11,13,41^. Moreover, our results suggest that global warming might generally give smaller species an edge in species interactions, resulting in asymmetries in species interactions^42,43^ that likely have significant consequences for community composition and ecosystem functions^7,44^.

Although previous research has often failed to detect acclimation in small organisms^18–20^, suggesting they might be at increased risk from GCC, our empirical and modeling results reveal that many small organisms, especially those at high latitudes, are indeed capable of rapid acclimation and because of this rapid acclimation, they have broad apparent thermal breadths.

To date, much of this acclimation has apparently gone undetected because of low heating rates in *CT*_*max*_ studies and delays in performance measurements that typify most experiments, or has been underestimated because most thermal plasticity studies ignore acclimation rates, which appear to be shorter for smaller organisms. Our results also suggest that researchers may be underestimating the plasticity of large organisms because many experiments do not provide sufficient time for them to fully acclimate to new temperatures. These results, coupled with many forecasts of GCC-induced extinctions not including behavioral or physiological (i.e., acclimation) plasticity to temperature^45,46^, suggest that some studies might have overestimated the risks of GCC to ectothermic animals. Recently, researchers came to similar conclusions for plants^7^. Such conclusions should not be taken as evidence that effects of GCC will not be catastrophic; however, it is at least a rare, albeit thin, silver lining in research on the effects of GCC on biodiversity. By providing a mechanistic understanding of acclimation based on geographic and species traits that are easily measured or inferred (i.e., latitude, ecto-vs endotherm, body size), we have provided an advance towards a framework for quantitatively predicting which ectothermic species and locations on the planet are most vulnerable to GCC, which should facilitate targeting limited conservation resources.

## Methods

### Data compilation

We tested our hypotheses about thermal acclimation and breadth using four independent datasets. The first and second datasets were compiled and described by Seebacher et al.^20^ and Gunderson and Stillman^23^ and offer 651 and 288 ARRs from studies with at least two acclimation temperatures, respectively. These datasets were reduced to 333 and 215 cases, respectively, with complete information and additional criteria applied (See SI Appendix, Table S1). The third database contains 2,445 thermal response curves of diverse performance traits ranging from feeding rate to body velocity^32,33^, to which we added data on acclimation temperatures and times (see below). The methods used to obtain and standardize these data are fully described in Dell et al.^32,33^. For some of our analyses, sample size was reduced to 128 of the 2,445 thermal responses (and 19 traits) for which there were non-monotonic performance curves (which are necessary to estimate optimal temperature, *T*_*opt*_) and acclimation temperature, location, and mass data. The fourth dataset consists of 1,040 estimates of *CT*_*maxs*_ for 251 amphibian species. Given that amphibians can show considerable variation in body mass from water uptake or dehydration, we used snout-vent length as our body size estimate for this dataset.

### Estimation of thermal response parameters

To calculate the parameters of each intraspecific thermal response in the Dell et al. dataset (i.e., *T*_opt_, curve height, and breadth), we used the *bbmle* package in R to fit unimodal functions to all non-monotonic temperature performance curves (those where the minimum tested temperature < *T*_opt_ < maximum tested temperature) with at least 5 points and assuming Gaussian distributed errors. We used Johnson-Lewin (Eq. 1)^32,33^ and Weibull (Eq. 2)^6^ functions to fit the thermal performance curves because both can fit asymmetrical curves without falling below zero on the y-axis (see SI Appendix for additional details). We eliminated fits where *T*_opt_ was outside of the range of temperatures tested. We calculated thermal breadths as the width of each thermal performance curve at 75% of the maximum height (*T*_opt_). Because breadth measurements that exceed the range of tested temperatures are unreliable, we excluded 13 cases where this occurred, resulting in a final sample size of 107.

### Overview of the mathematical model

Our model of thermal reaction norms (Fig. 4) assumed that: i) all organisms possess a common (identically broad) Gaussian (symmetric) or Weibull (asymmetric) thermal performance curve with a *T*_opt_ that depends on their latitude, ii) organisms acclimate to test temperatures that differ from their thermal optimum by translating (i.e., sliding) their thermal performance curves along the temperature axis, iii) the magnitude of acclimation depends on the organism’s acclimation rate and the acclimation duration up to some physiological limit of maximum acclimation, iv) acclimation rate scales allometrically with body mass and test temperature, and v) maximum acclimation depends linearly on absolute latitude. To generate predictions for the relationships among body size, latitude, acclimation, and performance breadth, we first simulated a pre-experiment laboratory acclimation period and then simulated an experiment in which 1,000 species were collected from various locations, acclimated to a given temperature in the laboratory for a fixed amount of time, and then performance was assessed across a temperature gradient. We assumed that organisms were allowed to acclimate to these experimental temperatures for a period of time that was shorter than the pre-experiment laboratory acclimation duration. Using the performance data simulated for each species at each temperature, we fitted Gaussian and Weibull thermal performance curves for each species using the *nls* function in the *stats* package in R. We then extracted parameters for *T*_opt_ and breadth (as the parameter *c*) from the Gaussian fits, and numerically computed these quantities, with breadth defined as the range over which organismal performance was ≥ 75% of peak performance, for the Weibull fits. We then analyzed these data with models that paralleled those used for the real dataset. See Methods in the SI Appendix for additional details.

### Statistical analyses

To test for effects of duration of time held at an acclimation temperature, we used the *lme* function in the *nlme* package of R statistical software to conduct a weighted mixed effects analysis (weighting by sample size and treating the study and species combination as a random effect) with *T*_opt_ or *CT*_max_ as the Gaussian response variables, habitat, trophic assignment (*T*_opt_ only), and life stage (*CT*_max_ only) as categorical moderators, and acclimation temperature, log acclimation duration, absolute value of latitude, and log body size as crossed continuous predictors (two- and three-way interactions only). To evaluate whether acclimation durations in our datasets were sufficient to acclimate large organisms, we repeated the acclimation duration analyses except we treated log acclimation duration as a response variable and excluded interactions.

To test the predictions of our mathematical model, we repeated the acclimation time analyses described above except we included all effect sizes where acclimation temperature data were available (see Tables S4-7 for details). For the Seebacher et al. analyses, our measure of acclimation strength was log(|1-Post-acclimation thermal sensitivity|+0.001)*-1. Post-acclimation thermal sensitivity was quantified in Seebacher et al.^20^ as a Q_10_ value where 1 indicates that physiological rates do not change with a change in acclimation temperatures. Thus, according to Seebacher et al.^20^ “the closer Q_10_ is to 1, the less affected animal physiology will be to a change in environmental temperature, meaning that animals will be more resilient to climate change”. Hence, because the direction of the change in a physiological rate will depend on the trait (e.g. swimming speed, metabolic rate, etc.), we took the absolute value of the deviation from 1. The log transformation was used to normalize the variable and multiplying by -1 resulted in more positive values intuitively indicating stronger acclimation. Results did not differ if we conducted analyses on both *in situ* and *ex situ* measurements (See SI Appendix, Table S5a) or on *in situ* whole body measurements only (See SI Appendix, Table S5b). Thus, we focus on analyses conducted on both *in situ* and *ex situ* measurements because it provided the larger sample size. For the Gunderson and Stillman analyses, ARR was the response variable and seasonality replaced latitude as a predictor. We then repeated these analyses on the breadth measurements from Dell et al.’s thermal performance curve dataset, again employing weighted mixed effect regression. To quantify the relationship between log body size and the time organisms were held at a test temperature before trait measurements were taken, we conducted a simple regression analysis using 1,480 of the 2,445 thermal response curves that had these data available. To validate our model of thermal acclimation and breadth, we treated amphibian species as the replicate (using the mean *CT*_*max*_ for each species) in the amphibian *CT*_*max*_ dataset, IUCN threat status as a binomial response variable, thermal safety margin, log body size, absolute value of latitude, log elevation, and log range size as crossed predictors, and a local estimate of the magnitude of climate change as a covariate (slope of the previous 50 years of maximum temperatures). To evaluate the contribution of acclimation plasticity to amphibian threat status, we analyzed the subset of amphibian *CT*_*max*_ data for which we also had ARR measurements, treating the magnitude of local climate change and log elevation as a covariates, and thermal safety margin, log body size, absolute value of latitude, and log ARR as crossed predictors. For all analyses, we chose not to include additional predictors that are included in some other acclimation studies, such as generation time and diel variation^6,40^. Specifically, we did not include generation time because it is highly collinear with log body size. We chose not to include diel variation because it is correlated with latitude and interacts with season. Because of this interaction and several studies not providing time of year of their collections, our sample sizes would have been further reduced if diel variation was included.

Where possible, we employed a multimodel inference approach (*dredge* and *model.avg* functions in the *MuMIn* package) to ensure we were not drawing conclusions based solely on one model. Multimodel inference compares all possible models using AIC and generates weighted coefficients and relative importance scores for predictors. We calculated conditional *R*^*^2^*^ values (variance explained by both fixed and random effects) for the best model where possible^47^, otherwise we calculated a *R*^*^2^*^ for the correlation between fitted and observed values. Analyses on *T*_opt_ and thermal breadth were conducted on both Johnson-Lewin and Weibull estimates of these parameters. For all analyses, log-likelihood ratio tests using the *Anova* function in the *car* package of R statistical software were used to calculate the probability values for each effect of the best performing model (i.e., lowest AIC). To display results of our regression models, we generated partial residual plots from the best model based on AIC using the *visreg* function in the *visreg* package of R statistical software. In all partial residual plots, continuous predictors are discretized strictly for the purposes of visually displaying statistical interactions (see SI Appendix for additional details). To ensure transparency, all datasets and code to reproduce the statistical analyses and figures are provided in a supplemental file.

### Data Accessibility

Data used for analyses in this manuscript can be found at http://biotraits.ucla.edu/, http://www.esapubs.org/archive/ecol/E094/108/, or in Database 1, which is an Excel file with 19 worksheets. One worksheet is the R code used to produce the figures and Tables in the paper. Nine of the remaining 18 worksheets are the datasets used for specific analyses in the paper. The remaining nine worksheets are the metadata that accompany each of the nine datasets.

## Acknowledgements

We thank S. Pawar, V. Savage, B. Garcia-Carreras, D. Kontopoulos, T. Smith, and an anonymous reviewer for helpful discussion or comments that improved this manuscript. This research was supported by grants from the National Science Foundation (EF-1241889, EID-1518681, EF-1241848), National Institutes of Health (R01GM109499, R01TW010286, F32AI112255), US Department of Agriculture (NRI 2006-01370, 2009-35102-0543), and US Environmental Protection Agency (CAREER 83518801) to J.R.R.

